# Ferrapoptosis: Discovery of a Hybrid Programmed Cell Death

**DOI:** 10.1101/2025.11.25.690347

**Authors:** Qiuyun Yang, Yang Chen, Yang Shen, Xiao Zhang, Xianliang Chen, Xinnan Zhao, Ruixuan Yuan, Manrui Li, Yang Xu, Xilong Lin, Yihan Sun, Shengyuan Zuo, Xinyang Liu, Zhuo Li, Ying Chen, Shichun Wang, Meili Lv, Lin Zhang, Weibo Liang, Xiameng Chen

## Abstract

Traditional taxonomies classify ferroptosis and apoptosis as distinct forms of regulated cell death. Here, we challenge this view by defining and validating a previously unrecognized neuronal death entity in traumatic brain injury (TBI), which we term “Ferrapoptosis”. This hybrid mode of death is characterized by the coexistence of ferroptotic and apoptotic molecular and ultrastructural features within the same neuron— ultrastructural alterations that cannot be fully characterized by any known cell death modalities. By integrating single-cell transcriptomics with multi-layered functional assays, we show that Ferrapoptosis dominates in severe injury and the acute phase, and is critically driven by mitochondrial oxidative stress. Over time, its predominance is gradually replaced by death modes in which either ferroptosis or apoptosis alone becomes the major pathway. Genome-wide CRISPR screening further identifies Smg7 as a key regulator that synchronously activates both death programs to drive the hybrid phenotype, whereas genetic or viral inhibition of Smg7 reduces Ferrapoptosis and promotes neurological recovery after TBI in mice. Our work systematically delineates a previously unrecognized form of cell death and its pathogenetic mechanisms, providing experimental evidence to refine cell death classification, and suggesting a conceptual strategy for treating complex diseases such as TBI by targeting shared regulatory nodes.

**HIGHLIGHTS:**

- Defined and validated a previously unrecognized cell death modality, termed “Ferrapoptosis”.
- Elucidated the ordered pattern of Ferrapoptosis across key stages of TBI pathogenesis.
- Revealed mitochondrial oxidative stress as the pivotal hub integrating ferroptosis and apoptosis.
- Identify Smg7 as a core driver of Ferrapoptosis and a promising neuroprotective target.

## INTRODCUTION

Traumatic brain injury (TBI) is a leading cause of death and disability worldwide, and survivors frequently develop long-term sequelae such as cognitive impairment, affective disturbances, and epilepsy [1]. Its pathological course begins with the immediate structural disruption caused by the primary mechanical insult and evolves into a secondary injury cascade that can persist from hours to years [2]. Progressive neuronal loss lies at the core of this process [3]. Accordingly, defining the precise modes of neuronal death after TBI and their molecular underpinnings has become a central question in the field. Over the past decade, the study of cell death has advanced beyond the traditional dichotomy of apoptosis between necrosis. The identification of ferroptosis, pyroptosis, necroptosis, and other forms of regulated cell death has greatly deepened our understanding of TBI pathophysiology [4]. Yet, beyond these established entities, it remains unclear whether the highly complex and heterogeneous microenvironment of TBI can drive additional, as-yet-unrecognized death programs. In the injured brain, excitotoxicity, ionic disequilibrium, inflammation, mitochondrial dysfunction, and reactive oxygen species (ROS) accumulation coexist, creating a milieu that favors the parallel activation of multiple death pathways. This leads us to a key question: in such a mixed stress environment, can cell death programs that were once viewed as strictly distinct breach their conventional boundaries and be engaged simultaneously within individual neurons?

In recent years, multiple studies have suggested that ferroptosis and apoptosis can be co-activated under diverse pathological conditions and engage in signaling crosstalk. At the molecular level, key regulatory nodes such as ROS bursts, mitochondrial dysfunction, BCL-2 family proteins, and p53 play pivotal roles in both pathways [4–6], indicating that they share components of the signaling machinery and providing a basis for inter-pathway communication. Recent work further shows that when the activity of GPX4, a core ferroptosis regulator, is inhibited, cells exhibit not only lipid peroxidation and other canonical ferroptotic features, but also apoptotic morphologies such as membrane blebbing [7]. Together, these findings suggest that ferroptosis and apoptosis are not fully independent, but under specific conditions have the potential for co-activation or cross-activation. However, several key questions remain: Is such co-activation or cross-activation sufficient to give rise to a bona fide novel cell death entity with characteristics of both types? What are its precise morphological phenotype, molecular signatures, and pathological dynamics? And how is it controlled by specific molecular regulators? To address these questions, we identify and validate in TBI a hybrid death modality—“Ferrapoptosis”—defined by the coexistence of canonical ferroptotic and apoptotic markers within single neurons. Through systematic analyses at both the phenotypic and mechanistic levels, we delineate the pathological characteristics of this death mode and further identify Smg7 as a key upstream regulator. These findings provide new experimental evidence for the complexity of cell death networks and may help refine and extend current classification schemes of regulated cell death.

## RESULTS

### TBI induces a hitherto uncharacterized form of neuronal death

By integrating published single-cell RNA-sequencing datasets of cortical neurons spanning the acute, subacute and chronic phases after TBI, we identified a neuronal subpopulation with concomitantly elevated ferroptosis and apoptosis pathway activity (“Double-High” neurons), which was clearly segregated from neurons with activation of only one pathway (Fig. 1A). Temporal quantification of pathway activity showed that Double-High neurons dominated the acute phase, whereas high-apoptosis and high-ferroptosis neurons became the major populations in the subacute and chronic phases, respectively (Fig. 1B-1C). These data suggest that ferroptotic and apoptotic features can coexist within single neurons and that activation of these two death programs after TBI is temporally organized. We next used gene set enrichment analysis (GSEA) to assess the functional state of Double-High neurons. Compared with neurons with high ferroptosis activity alone, Double-High neurons showed significant enrichment of apoptosis- and oxidative stress–related pathways; compared with neurons with high apoptosis activity alone, they displayed stronger immune activation signatures (Fig. 1D). These findings indicate that Double-High neuronal death is not a simple sum of two independent death programs.

**Figure 1.**
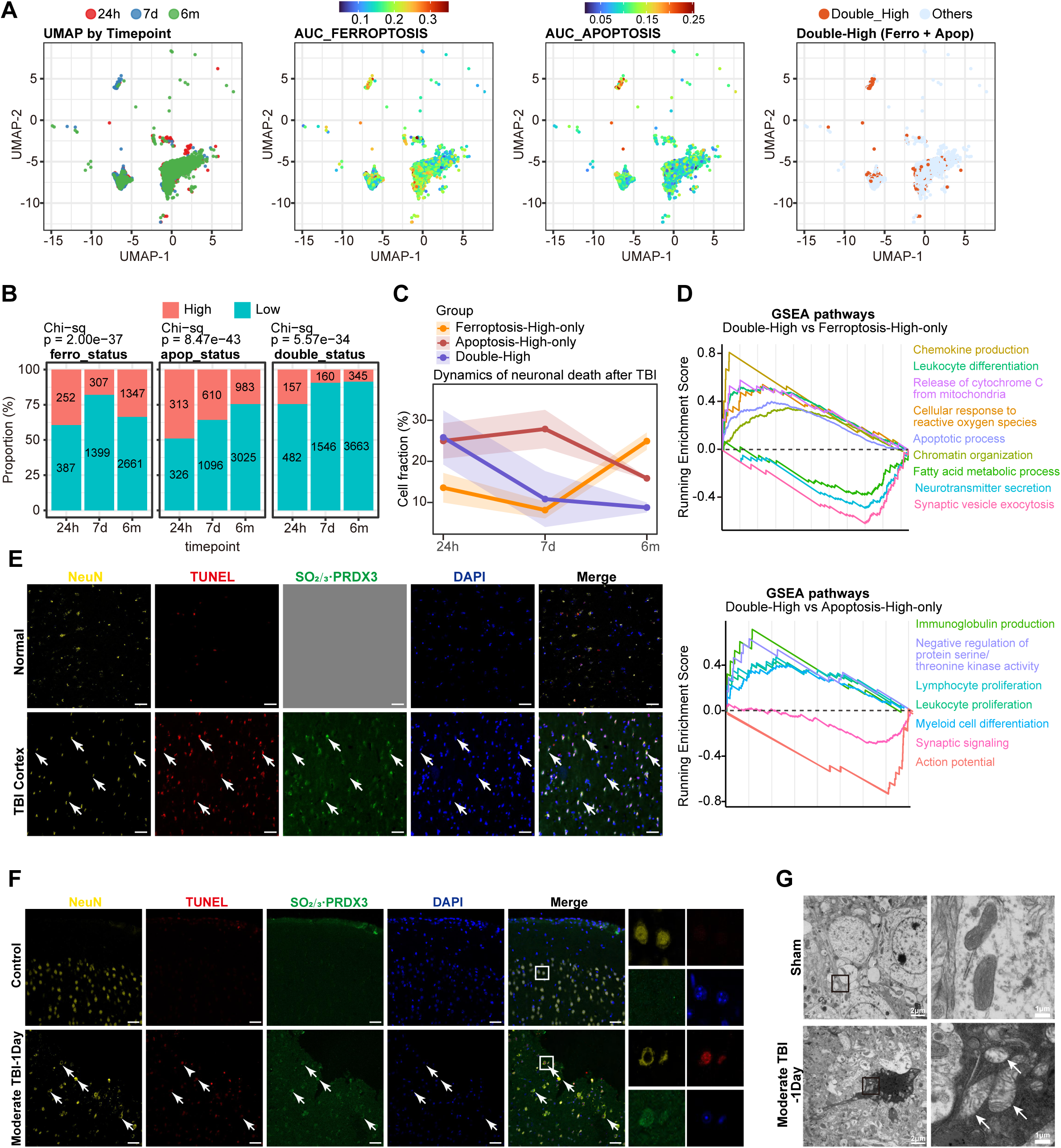
Identification of a hybrid neuronal death type “Ferrapoptosis” after TBI. **(A)** UMAP embedding of single-nucleus transcriptomes from cortical neurons at 24 h, 7 d and 6 months after TBI, colored by time point (left), ferroptosis gene-set score, apoptosis gene-set score, and by Double-High neurons (simultaneously high ferroptosis and apoptosis scores) versus all other neurons (right). **(B)** Bar plots showing the proportions of ferroptosis-High versus -Low, apoptosis-High versus -Low, and Double-High versus Double-Low neurons at each time point (Chi-square tests, P values indicated). **(C)** Line plots depicting the temporal dynamics of ferroptosis-High-only, apoptosis-High-only and Double-High neurons after TBI. **(D)** GSEA plots of representative pathways enriched in Double-High neurons compared with ferroptosis-High-only (up) or apoptosis-High-only (down) neurons. **(E-F)** Representative confocal images of human cortical sections from normal and TBI brains **(E)** and mouse cortical sections from sham and moderate TBI mice 1 day after injury **(F)** stained for NeuN (neurons, yellow), TUNEL (apoptosis, red), oxidized PRDX3-SO□/□ (mitochondrial oxidative stress/ferroptosis marker, green) and DAPI (nuclei, blue), scale bars=50μm. **(G)** Transmission electron micrographs of cortical neurons from sham and moderate TBI mice (1 day). Sham neurons display normal nuclear and mitochondrial morphology, whereas TBI neurons show apoptotic changes (chromatin condensation and margination) together with swollen mitochondria with reduced cristae indicated by arrows, Scale bars, as indicated.

To validate the coexistence of ferroptosis and apoptosis in neurons, we performed NeuN (neurons), TUNEL (apoptosis) and PRDX3-SO□/□ (ferroptosis) [8] triple immunofluorescence staining in human TBI specimens. Numerous triple-positive cells were detected in the injured cortex (Fig. 1E), pointing to a previously unrecognized dual-phenotype death mode. This pattern was recapitulated in a mouse moderate TBI model, where abundant triple-positive neurons were observed 24 h after injury (Fig. 1F). At the ultrastructural level, transmission electron microscopy further demonstrated the coexistence, within single neurons, of apoptotic features (chromatin condensation and margination) together with ferroptotic features (swollen mitochondria with reduced cristae) (Fig. 1G), revealing a unique ultrastructural profile that defies classification into any previously defined cell death modality. These morphological data substantiate the presence of a ferroptosis–apoptosis hybrid form of neuronal death, hereafter referred to as “Ferrapoptosis.”

### Pathological features of Ferrapoptosis across injury severity and time

To investigate how Ferrapoptosis relates to TBI severity and post-injury time, we established mouse TBI models encompassing different injury grades and time points (Fig. 2A). Successful TBI induction in mice was verified by neurological assessment using the modified Neurological Severity Score (mNSS) (Fig. 2B). At a given time point, the number of TUNEL□/PRDX3-SO□/□□ double-positive neurons increased markedly with escalating injury severity (Fig. 2C): abundant triple-positive neurons were observed in the cortex after severe TBI, an intermediate number after moderate TBI, and only scattered cells after mild TBI. These dual-phenotype signals were largely confined to the ipsilateral hemisphere, with only rare cells detected contralaterally (Fig. S1), indicating that Ferrapoptosis is tightly linked to focal lesion severity .

**Figure 2.**
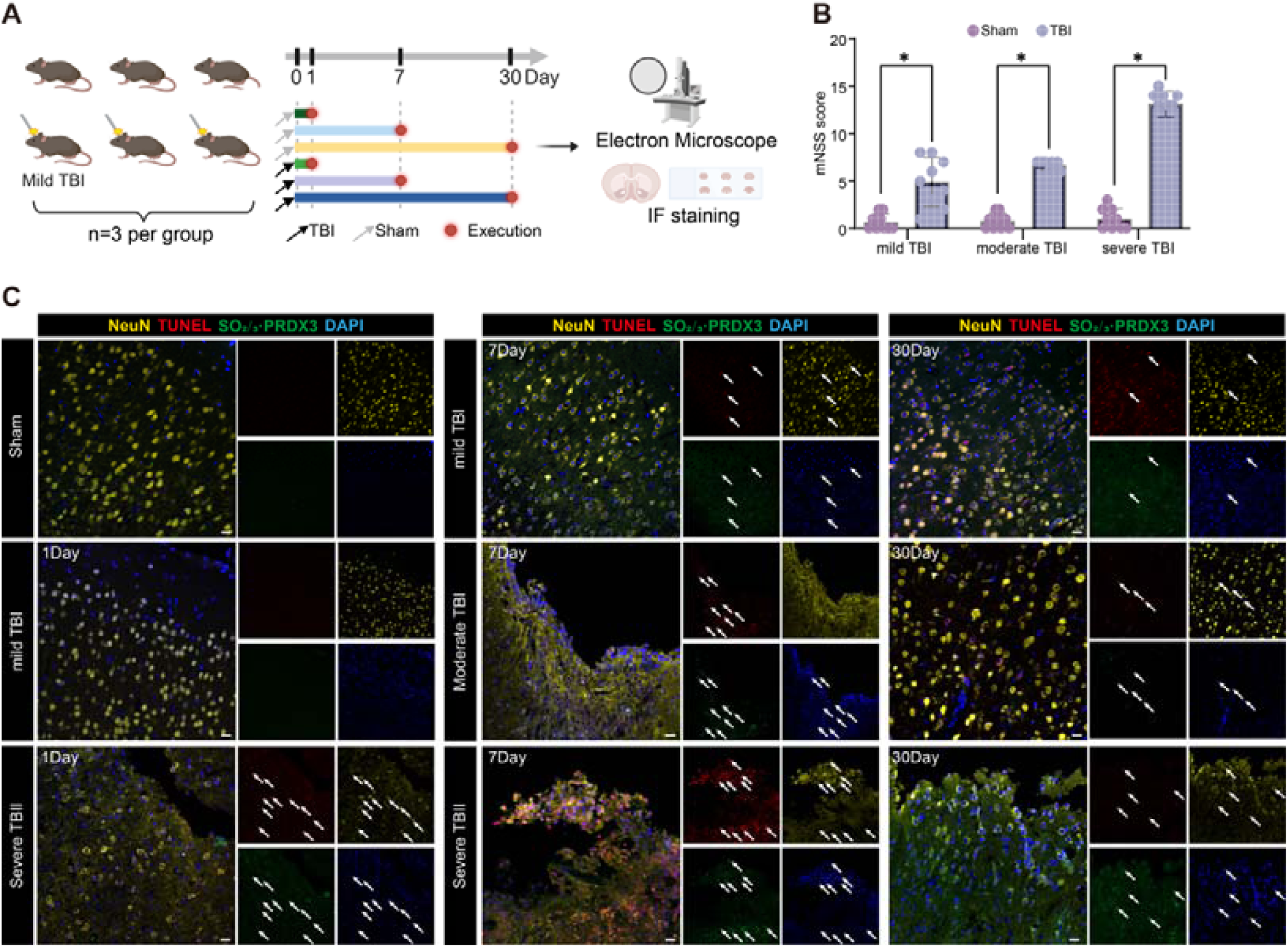
Ferrapoptosis varies with TBI severity and evolves over time. **(A)** Schematic of the in vivo experimental design. Mild, moderate and severe TBI were induced by controlled cortical impact; mice were evaluated at 1, 7 and 30 days after injury (n = 3 per group) and brains were processed for immunofluorescence and electron microscopy. **(B)** mNSS in mice subjected to mild, moderate, or severe TBI compared with sham. Data are presented as mean ± SEM (n = 9 per group). Statistical significance was analyzed by unpaired two-tailed Student’s t-test between Sham and TBI within each injury grade; *P < 0.05. **(C)** Representative confocal images of ipsilateral cortex and hippocampus from sham, mild, moderate and severe TBI mice at 1, 7 and 30 days post-injury stained for NeuN (yellow), TUNEL (red), PRDX3-SO□/□ (green) and DAPI (blue), arrows indicate triple-positive neurons, scale bars=50μm.

Across time, double-positive neurons generally peaked at day 1 after injury in most brain regions and, although partially reduced by day 30, remained significantly higher than in sham-operated mice (Fig. 2C). This dynamic pattern suggests that Ferrapoptosis is most prominent in the acute phase and only partially resolves in the chronic phase, consistent with the temporal death signatures revealed by our single-cell transcriptomic analyses (Fig. 1B-1C). Together, these data indicate that the pathological engagement of Ferrapoptosis is jointly shaped by both TBI severity and time after injury.

### Selective induction of ferroptosis or apoptosis is sufficient to perturb the other death program

To dissect crosstalk between apoptosis and ferroptosis, we treated HT22 cells with staurosporine (STS; apoptosis inducer), Erastin (ferroptosis inducer), sodium citrate (reported to concomitantly trigger ferroptosis and apoptosis [9]), or mechanical scratch injury to mimic TBI in vitro [10], and assessed morphology and viability 12 h later. All four insults reduced cell viability and increased cell death relative to control (Fig. 3A-3B). Morphologically, STS induced classical apoptotic features such as membrane blebbing, whereas Erastin caused cell swelling typical of ferroptosis. By contrast, in the scratch and sodium citrate groups we observed cells in the same field displaying apoptotic-like membrane blebs together with ferroptosis-like somatic swelling, suggesting the presence of non-classical death responses.

**Figure 3.**
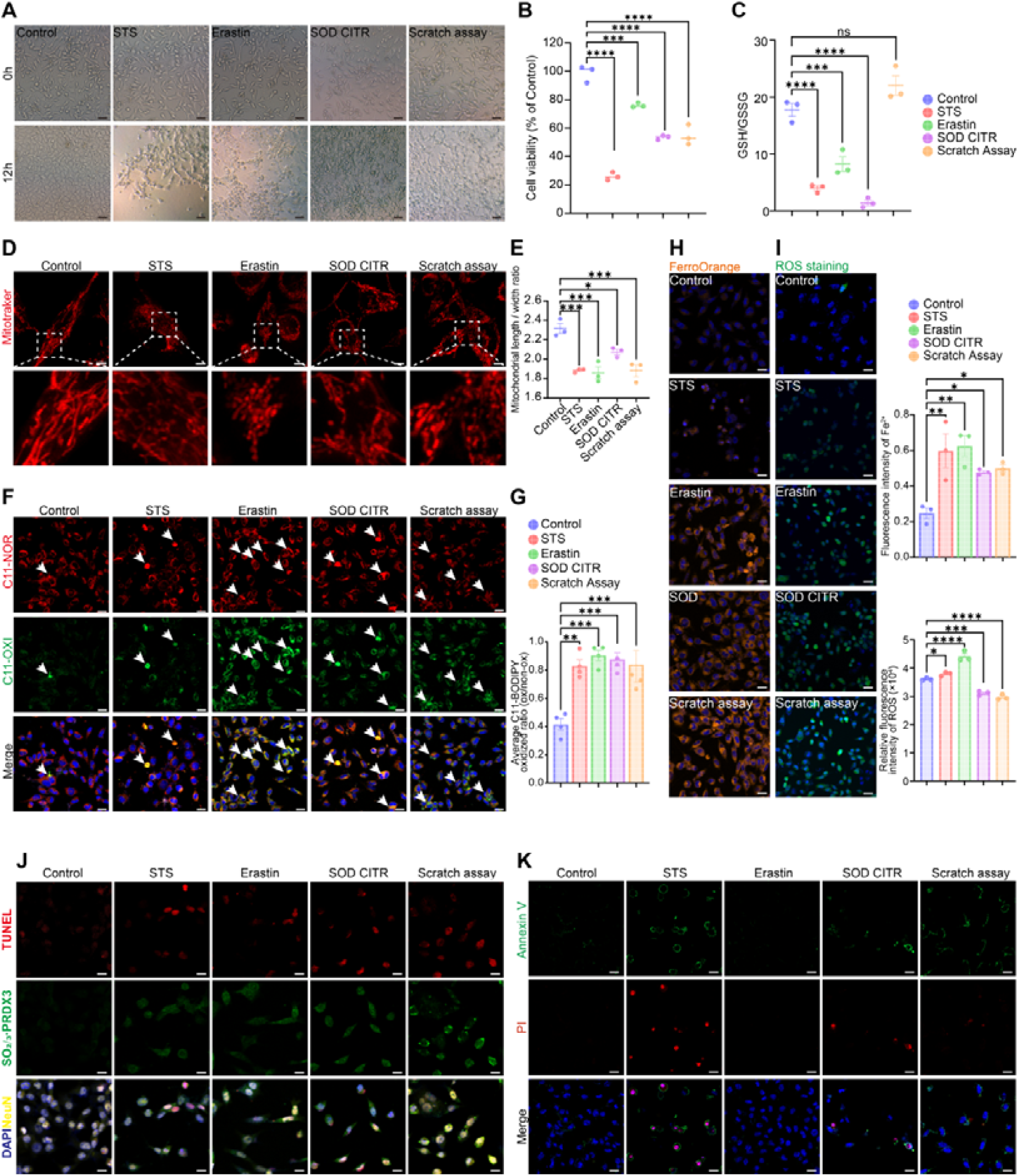
Distinct death stimuli in vitro converge on mitochondrial damage and ferroptosis-apoptosis crosstalk. **(A)** Phase-contrast images of HT22 cells at 0 and 12 h after treatment with vehicle (Control), staurosporine (STS), erastin, sodium citrate (SOD CITR) or scratch injury, scale bars=20μm. **(B)** Cell viability of HT22 cells measured by CCK-8 assay 12 h after treatment with vehicle (Control), STS, Erastin, sodium citrate (SOD CITR) or scratch injury. Data are presented as mean ± SEM (dots represent independent cultures, n = 3 per group). Statistical analysis was performed by one-way ANOVA followed by Tukey’s post hoc test; ***P < 0.001, ****P < 0.0001. **(C)** Intracellular redox status assessed by the GSH/GSSG ratio in HT22 cells under the same treatment conditions as in (B). Data are presented as mean ± SEM (n = 3 per group). Group differences were analyzed by one-way ANOVA with Tukey’s post hoc test; ns, not significant; ***P < 0.001, ****P < 0.0001. **(D)** Mitotracker staining of HT22 cells after treatment to visualize mitochondrial morphology, scale bars=5μm. **(E)** Quantification of mitochondrial length/width ratio from Mitotracker images in **(D)**. Data are presented as mean ± SEM (n = 3 per group). Statistical analysis was performed by one-way ANOVA followed by Tukey’s post hoc test; **P < 0.01, ***P < 0.001. **(F)** C11-BODIPY 581/591 staining of HT22 cells under the indicated treatments to assess lipid peroxidation (C11-NOR, reduced form; C11-OXI, oxidized form). Nuclei are counterstained with Hoechst, scale bars=20μm. **(G)** Quantification of C11-BODIPY oxidized fluorescence from images in (F). Data are mean ± SEM (n = 3 independent cultures per group). Group differences were analyzed by one-way ANOVA with Tukey’s post hoc test; **P < 0.01, ***P < 0.001. **(H-I)** FerroOrange staining **(H)** and ROS staining **(I)** of HT22 cells to detect intracellular Fe²□ after the indicated treatments. Nuclei are counterstained with Hoechst, scale bars=20μm. Representative images (left) and quantification of mean fluorescence intensity (right). Data are mean ± SEM (n = 3 per group). One-way ANOVA with Tukey’s post hoc test was used; *P < 0.05, **P < 0.01, ***P < 0.001, ****P < 0.0001. **(J)** Immunofluorescence staining of HT22 cells for TUNEL, PRDX3-SO□/□ and DAPI to evaluate apoptotic and ferroptotic markers and their co-localization under the indicated treatments, scale bars=20μm. **(K)** Annexin V–FITC (green)/PI (red) staining of HT22 cells with Hoechst nuclear counterstaining under the indicated treatments to visualize early apoptotic (Annexin V□/PI□) and late apoptotic/necrotic (Annexin V□/PI□) cells, scale bars=20μm.

At the biochemical level, STS-induced apoptosis was accompanied by a decrease in the GSH/GSSG ratio (Fig. 3C), indicating that a “pure” apoptotic stimulus can also engage ferroptosis-related metabolic changes. Notably, in the scratch model, GSH levels were paradoxically increased, which we speculate reflects activation of endogenous antioxidant defenses in response to intense oxidative stress. Similar to Erastin-induced ferroptosis, STS treatment reduced mitochondrial aspect ratio (Fig. 3D-3E) and increased lipid peroxidation (Fig. 3F-3G), ferrous iron accumulation (Fig. 3H), and ROS production (Fig. 3I). Together, these data show that even a single apoptotic trigger such as STS can substantially perturb iron homeostasis and ferroptotic processes, reinforcing the notion of tight bidirectional crosstalk between the two death pathways.

Consistent with our in vivo findings, NeuN□/TUNEL□/PRDX3-SO□/□□ triple-positive Ferrapoptosis neurons were also detected in the scratch and sodium citrate groups in vitro (Fig. 3J). Apoptotic readouts based on Annexin V/PI staining confirmed robust apoptosis in the STS group, and all non-control groups displayed a loss of mitochondrial membrane potential comparable to that seen with STS (Fig. 3K), as evidenced by both the JC-1 red/green shift and quantitative aggregate/monomer ratios (Fig. S2). Collectively, these results demonstrate that selective induction of either ferroptosis or apoptosis is sufficient to disturb the other program, and that such bidirectional crosstalk likely forms a key mechanistic basis for the emergence of the Ferrapoptosis hybrid death mode.

### Mitochondrial oxidative stress is a key driver of Ferrapoptosis

Building on the above findings, we noted that Ferrapoptosis, ferroptosis and apoptosis are all accompanied by abnormal mitochondrial morphology, loss of mitochondrial membrane potential, and accumulation of intracellular lipid peroxides and ROS. This suggested that Ferrapoptosis may be driven by exacerbated mitochondrial oxidative stress. To test this, we further examined the role of mitochondria in this hybrid death mode. Treatment with the mitochondria-targeted antioxidant MitoQ markedly attenuated both apoptotic (TUNEL) and ferroptotic (PRDX3-SO□/□) signals in STS-, Erastin-, sodium citrate– and scratch-injured cultures, and reduced the proportion of TUNEL□/PRDX3□ ferrapoptotic neurons (Fig. 4A-4B). Consistently, flow cytometry showed that MitoQ pretreatment significantly decreased Annexin V and C11-BODIPY mean fluorescence intensity in the treated groups (Fig. 4C-4F). Together, these results indicate that mitochondrial oxidative stress is a critical driving force for Ferrapoptosis, and that its inhibition can effectively prevent downstream cross-activation of death signals, establishing mitochondria as a central regulator of this hybrid cell death program.

**Figure 4.**
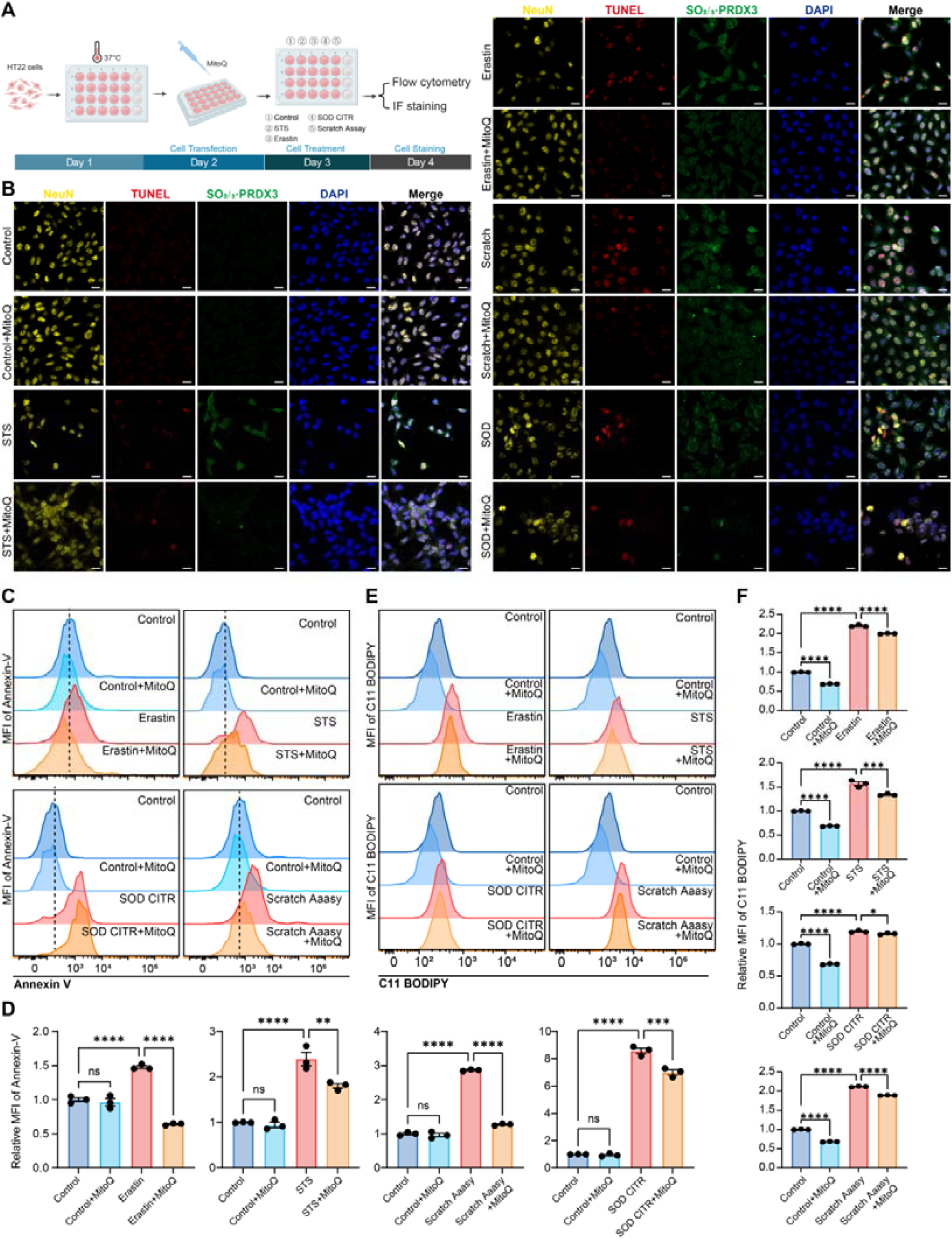
Mitochondria-targeted antioxidant MitoQ attenuates Ferrapoptosis-like phenotypes. **(A)** Schematic of the in vitro HT22 experimental workflow. **(B)** Representative IF images of HT22 cells stained for NeuN (yellow), TUNEL (red), PRDX3-SO□/□ (green) and DAPI (blue) under the indicated conditions, with or without MitoQ pretreatment, scale bars=20μm. **(C)** Flow-cytometry histograms of Annexin V-FITC fluorescence showing that MitoQ decreases apoptosis under erastin, STS, SOD CITR and scratch treatment. **(D)** Quantification of Annexin V mean fluorescence intensity across conditions. **(E)** Flow-cytometry histograms of C11-BODIPY fluorescence demonstrating that MitoQ lowers lipid peroxidation in the presence of different death stimuli. **(F)** Quantification of C11-BODIPY MFI showing reduced lipid peroxidation with MitoQ. Data are mean ± SEM from independent experiments; P values are shown.

### Smg7 is a key regulator driving Ferrapoptosis

To dissect the molecular regulation of Ferrapoptosis, we first leveraged two genome-wide CRISPR–Cas9 knockout screening datasets in mice and identified 2,610 candidate genes linked to ferroptosis and 23 linked to apoptosis. Given that Ferrapoptosis integrates features of both death programs, we intersected these gene sets and obtained 10 high-priority candidates that may co-regulate both pathways: *Acsl4*, *Phf12*, *4931417G12Rik*, *Tnfrsf1b*, *Tnfrsf1a*, *Bid*, *Casp8*, *Tc2n*, *Ces2a* and *Smg7* (Fig. 5A). Literature mining suggested that Smg7 is closely associated with cellular oxidative stress responses [11–13], pointing to a potential role in the core pathological axis of Ferrapoptosis—mitochondrial oxidative stress.

**Figure 5.**
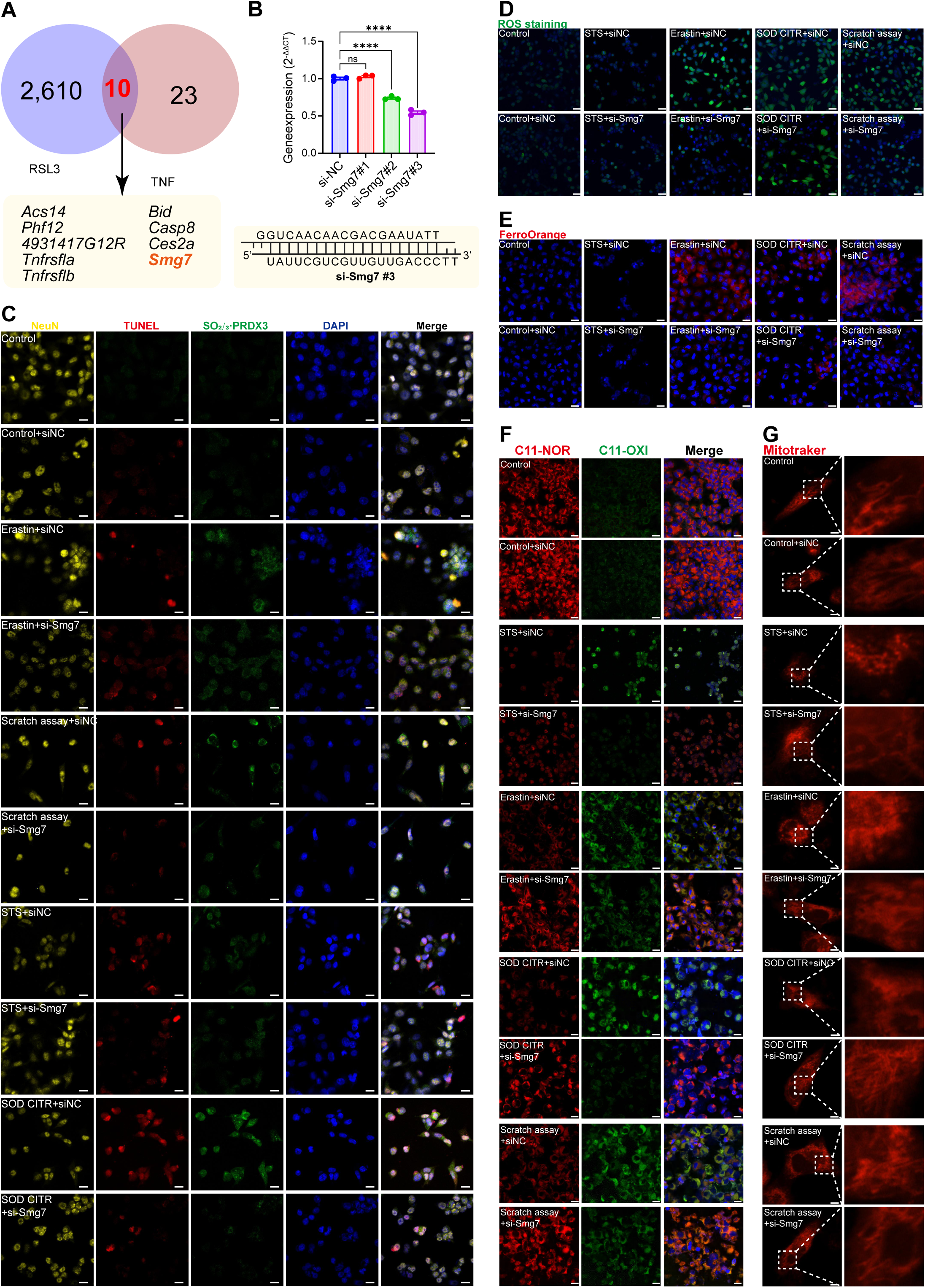
Genome-wide screening identifies Smg7 as a regulator of Ferrapoptosis. **(A)** Venn diagram of CRISPR-Cas9 loss-of-function screens performed under RSL3-induced ferroptosis and TNF-induced apoptosis, revealing 10 overlapping candidate genes that modulate both death pathways. **(B)** qPCR validation of Smg7 knockdown efficiency in HT22 cells transfected with three independent siRNAs and a negative control (si-NC); si-Smg7#3 is selected for subsequent experiments (sequence shown below). **(C)** Immunofluorescence staining for NeuN (yellow), TUNEL (red), PRDX3-SO□/□ (green) and DAPI (blue) in HT22 cells under the indicated treatment conditions with or without si-Smg7, scale bars=20μm. **(D)** General ROS staining of HT22 cells under the indicated treatment conditions with or without si-Smg7; nuclei are counterstained with Hoechst, scale bars=20μm. **(E)** FerroOrange staining of intracellular Fe²□ in HT22 cells under the indicated conditions with or without si-Smg7; nuclei are counterstained with Hoechst, scale bars=20μm. **(F)** C11-BODIPY 581/591 staining of HT22 cells under the indicated treatments with or without si-Smg7 to assess lipid peroxidation (C11-NOR, reduced form, red; C11-OXI, oxidized form, green); nuclei are counterstained with Hoechst, scale bars=20μm. **(G)** Mitotracker Red staining of HT22 cells under the indicated conditions with or without si-Smg7 to visualize mitochondrial morphology, scale bars=5μm.

To test the involvement of Smg7 in Ferrapoptosis, we performed siRNA-mediated knockdown (Fig. 5B). Compared with the si-NC control, si-Smg7 markedly reduced the proportion of TUNEL□/oxidized PRDX3□ double-positive cells (Fig. 5C), indicating that Smg7 silencing effectively suppresses neuronal death co-mediated by ferroptosis and apoptosis. In parallel, si-Smg7 significantly lowered intracellular ROS (Fig. 5D), ferrous iron levels (Fig. 5E) and lipid peroxidation (Fig. 5F), suggesting that Smg7 knockdown alleviates oxidative stress and halts ferroptotic progression. Mitotracker staining further showed that si-Smg7 helps restore an elongated, spindle-shaped mitochondrial morphology (Fig. 5G), supporting a role for Smg7 in maintaining mitochondrial structural integrity. Together, these data indicate that Smg7 promotes ferroptosis, apoptosis and Ferrapoptosis by enhancing mitochondrial oxidative stress and lipid peroxidation.

At the level of apoptotic readouts, si-Smg7 restored mitochondrial membrane potential (Fig. 6A-B) and significantly reduced the percentage of PI□/Annexin V□ cells (Fig. 6C-D), further confirming its impact on apoptosis. Conversely, Smg7 overexpression exacerbated ferroptotic, apoptotic and ferrapoptotic phenotypes (Fig. 6E-6F, Supplementary File 1).

**Figure 6.**
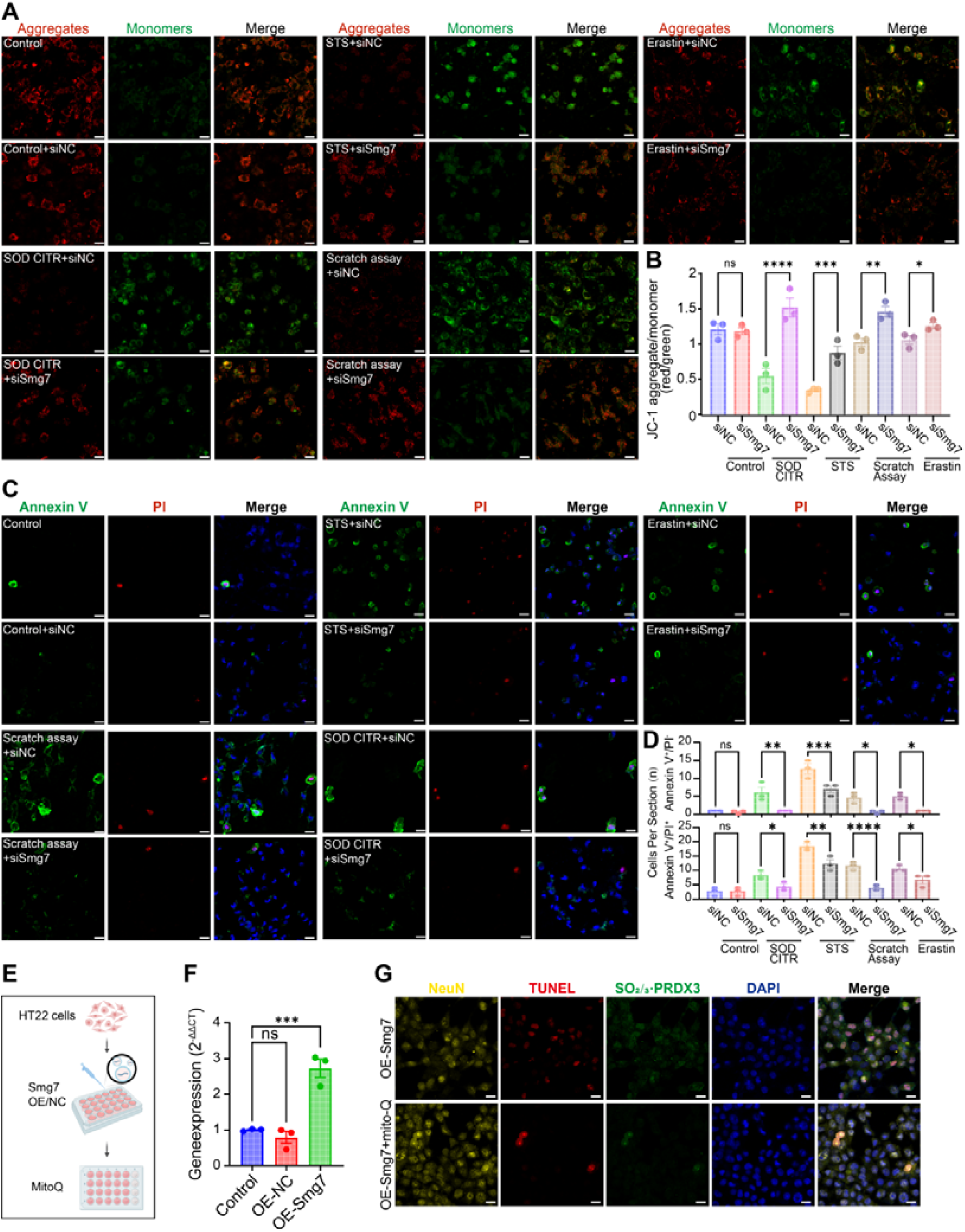
Smg7 modulates mitochondrial membrane potential and apoptotic execution. **(A)** JC-1 staining of HT22 cells under the indicated treatment conditions with or without si-Smg7 to assess mitochondrial membrane potential (JC-1 aggregates, red; monomers, green); nuclei are counterstained with Hoechst, scale bars=20μm. **(B)** Quantification of the JC-1 red/green fluorescence ratio from images in (A). Data are presented as mean ± SEM (n = 3 per group). Group differences were analyzed by one-way ANOVA followed by Tukey’s post hoc test; ns, not significant; *P < 0.05, **P < 0.01, ***P < 0.001. **(C)** Annexin V–FITC/PI staining of HT22 cells under the indicated treatment conditions with or without si-Smg7; nuclei are counterstained with Hoechst, scale bars=20μm. **(D)** Quantification of Annexin V□ and Annexin V□/PI□ cells per field from the conditions shown in (C). Data are presented as mean ± SEM (n = 3 independent cultures per group) and were analyzed by one-way ANOVA with Tukey’s post hoc test; ns, not significant; *P < 0.05, **P < 0.01. **(E)** Schematic of the experimental workflow for Smg7 overexpression (OE-Smg7) in HT22 cells and subsequent MitoQ treatment. **(F)** Relative Smg7 mRNA expression measured by qPCR in control, empty-vector (OE-NC) and OE-Smg7 HT22 cells. Data are shown as mean ± SEM (n = 3 independent experiments) and were analyzed by one-way ANOVA with Tukey’s post hoc test; ns, not significant; ***P < 0.001. **(G)** Immunofluorescence staining for NeuN (yellow), TUNEL (red), PRDX3-SO□/□ (green) and Hoechst (blue) in HT22 cells expressing OE-Smg7 with or without MitoQ, under the indicated conditions, scale bars=20μm.

Collectively, these findings identify Smg7 as a key molecular regulator of Ferrapoptosis. By amplifying mitochondrial oxidative stress, promoting lipid peroxidation and facilitating ferrous iron accumulation, Smg7 coordinately activates ferroptotic and apoptotic signaling, thereby driving the progression of this hybrid cell death mode.

### Smg7 drives Ferrapoptosis through transcriptional reprogramming and shapes neurological recovery

To elucidate how Smg7 regulates Ferrapoptosis, we performed transcriptome profiling of scratch-injured HT22 cells treated with si-NC or si-Smg7 (Fig. 7A-7B). Smg7 knockdown partially reversed scratch-induced changes in key regulators of ferroptosis and apoptosis (Fig. 7C-7D). In the ferroptosis pathway, expression of *Hmox1*, *Ncoa4* and *Ftl1* was markedly reduced after si-Smg7 treatment. In the apoptosis pathway, the pro-apoptotic factors *Apaf1* and *Casp9* were downregulated, whereas the anti-apoptotic gene *Bcl2* was upregulated. Notably, in contrast to classical pharmacological models, *Acsl4*, *Tfrc* and *Bid* were downregulated in the scratch model, suggesting mechanistic heterogeneity in how different injury paradigms engage cell death programs. Gene Ontology enrichment analysis further showed that si-Smg7 robustly activated pathways related to intrinsic apoptosis inhibition, mitochondrial regulation and synaptic repair (Fig. 7E), indicating that Smg7 suppression shifts neurons from a pro-death state toward a pro-survival, synaptic remodeling–oriented state. Follow-up in vivo validation confirmed that the expression trends of all genes except *Hmox1* were consistent with the RNA-seq results (Fig. 7F, Supplementary File 2). Together, these data indicate that Smg7 drives Ferrapoptosis by coordinately remodeling the expression of multiple key genes in the ferroptotic and apoptotic cascades.

**Figure 7.**
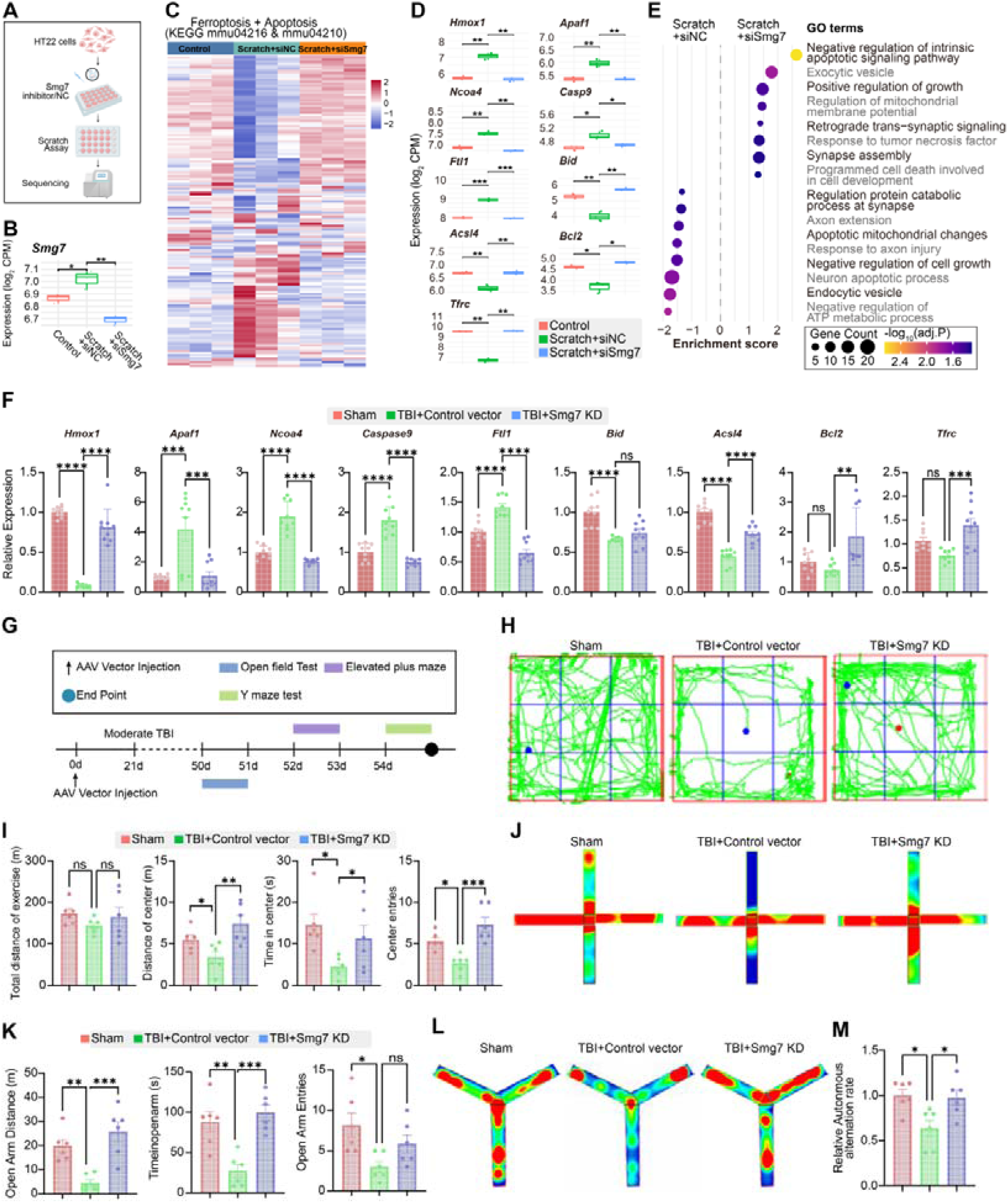
Smg7-dependent transcriptional reprogramming <u>drives</u> Ferrapoptosis and impairs functional recovery after TBI. Schematic of the bulk RNA-seq workflow in HT22 cells subjected to scratch injury with or without si-Smg7. **(B)** Smg7 mRNA expression levels in HT22 cells across the indicated groups. Data are presented as mean ± SEM (n = 3 per group). Group differences were analyzed by one-way ANOVA followed by Tukey’s post hoc test; ns, not significant; *P < 0.05, **P < 0.01, ***P < 0.001. **(C)** Heatmap of genes belonging to the KEGG ferroptosis (mmu04216) and apoptosis (mmu04210) pathways across the indicated RNA-seq groups. Values represent row-scaled (Z-score) normalized expression. **(D)** Box plots of representative ferroptosis- and apoptosis-related genes from the dataset shown in (C) across the indicated RNA-seq groups. Statistical significance was determined using DESeq2 Wald tests with Benjamini–Hochberg correction; *P < 0.05, **P < 0.01, ***P < 0.001, ****P < 0.0001. **(E)** GO enrichment analysis of differentially expressed genes between scratch-injured cells with or without Smg7 knockdown. Dot size indicates gene counts and color denotes -log10(FDR). **(F)** qPCR analysis of selected ferroptosis- and apoptosis-related genes in cortical tissue from the indicated mouse groups. Data are shown as mean ± SEM (n = 3 per group) and were analyzed by one-way ANOVA with Tukey’s post hoc test; *P < 0.05, **P < 0.01, ***P < 0.001, ****P < 0.0001. **(G)** Experimental timeline illustrating AAV vector injection, induction of moderate TBI, subsequent behavioral testing. **(H)** Representative open-field tracking plots from the three groups 30 days after injury. **(I)** Quantification of open-field parameters corresponding to (G), including total distance traveled, distance traveled in the center zone, time spent in the center and number of center entries. Data are presented as mean ± SEM (n = 6 mice per group). Statistical comparisons were performed by one-way ANOVA followed by Tukey’s post hoc test; ns, not significant; *P < 0.05, **P < 0.01, ***P < 0.001. **(J)** Representative tracking plots in the elevated plus maze for the same three groups. **(K)** Quantification of elevated plus maze parameters corresponding to (I). Data are mean ± SEM (n = 6 mice per group) and were analyzed by one-way ANOVA with Tukey’s post hoc test; ns, not significant; *P < 0.05, **P < 0.01, ***P < 0.001. **(L)** Representative tracking plots in the Y-maze spontaneous alternation test for the indicated mouse groups. **(M)** Relative spontaneous alternation rate in the Y-maze. Data are presented as mean ± SEM (n = 6 mice per group) and were analyzed by one-way ANOVA followed by Tukey’s post hoc test; *P < 0.05.

We next asked whether Smg7 modulation impacts functional recovery after TBI. One month after injury, we assessed motor and cognitive performance in mice (7G). In the open-field test, total distance traveled did not differ significantly among groups, but si-Smg7–treated mice displayed increased locomotion, time spent and entries into the center zone compared with TBI+si-NC mice (Fig. 7H-7I), indicating an improvement in anxiety-like behavior. Consistently, the elevated plus maze showed that si-Smg7 alleviated TBI-induced anxiety and enhanced exploratory behavior (Fig. 7J-7K). In the Y-maze, si-Smg7 significantly increased spontaneous alternation, reflecting a recovery of spatial working memory (Fig. 7L-7M). Collectively, these findings demonstrate that targeting Smg7 not only suppresses Ferrapoptosis at the cellular level, but also promotes long-term neurobehavioral recovery after TBI.

## DISCUSSION

Over the past decade, the taxonomy of cell death has expanded from the classical dichotomy of apoptosis versus necrosis into a complex spectrum that includes pyroptosis, ferroptosis, cuproptosis and other forms of regulated cell death, each with distinct ultrastructural features, signaling pathways and physiological roles [14, 15]. Here, we identify and define a previously unrecognized death modality, “Ferrapoptosis,” and provide convergent evidence for its existence across human TBI specimens, animal models and single-cell transcriptomic datasets. This concept is conceptually important: ferroptosis and apoptosis have traditionally been viewed as fundamentally distinct entities [14], yet our data demonstrate that they are not strictly mutually exclusive and can coexist within the same neuron. The discovery of this hybrid death program is expected to inform future refinements of the current classification framework of regulated cell death.

We further show that the activation of ferroptosis and apoptosis in the TBI milieu follows a distinct temporal trajectory. The large number of ferrapoptotic neurons observed in the acute phase likely arises from the combination of intense primary mechanical injury, burst of free radicals, calcium overload and acute inflammation [16–18], all of which favor the simultaneous engagement of multiple death programs. Mechanical impact and microhemorrhage rapidly increase free iron and trigger ROS surges, promoting lipid peroxidation and membrane damage; in parallel, glutamate excitotoxicity and mitochondrial dysfunction activate the canonical apoptotic pathway [19]. As injury progresses into the subacute phase, acute responses begin to subside and reparative processes are initiated. By this time, cells that sustained irreversible damage during the initial insult might have largely completed their death programs. The remaining neurons are exposed to more moderate, chronic stress, which is more likely to preferentially engage a single death pathway adapted to persistent low-level insult, with classical apoptosis progressively becoming dominant. At later stages, the microenvironment becomes relatively stable, but ongoing oxidative stress and iron accumulation emerge as key drivers. As pro-apoptotic signals wane, iron dysregulation and sustained lipid peroxidation may render ferroptosis the predominant mode of cell death in the chronic phase [20]. We also observed that the proportion of Ferrapoptosis closely tracks TBI severity. Severe injury typically causes abrupt disruption of membrane integrity, profound energetic collapse and massive oxidative stress [21]. Such an “explosive” injury pattern may concurrently and strongly activate multiple death cascades, accounting for the marked increase in ferrapoptotic neurons in this setting. By contrast, mild-to-moderate TBI induces relatively milder stress, allowing cells to maintain partial energy metabolism and membrane integrity for a period of time. This might create a window for more orderly engagement of signaling pathways, favoring either highly programmed apoptosis or a slower progression toward ferroptosis under sustained sublethal oxidative stress. These observations have potential implications for stage- and severity-specific interventions in TBI, and suggest that tailored therapeutic strategies targeting distinct death pathways at different time windows may be required to achieve truly precise treatment.

In this study, we also identify Smg7 through genome-wide knockout datasets and demonstrate its central role in driving Ferrapoptosis using functional assays. Smg7 is a core component of the nonsense-mediated mRNA decay (NMD) machinery, classically known for safeguarding mRNA quality. Previous work has shown that Smg7 can potentiate TNF-α receptor-mediated extrinsic apoptosis [22] and interact with p53 to regulate intrinsic apoptosis in the DNA damage response [12], yet its involvement in ferroptosis has not been reported. Here, we demonstrate for the first time that Smg7 functions as an upstream hub that coordinately regulates mitochondrial apoptosis and ferroptosis. On the one hand, Smg7 may facilitate intrinsic apoptosis by modulating apoptosome assembly (*Apaf1*-*Casp9*) [23] and the balance within the Bcl-2 family (*Bid*/*Bcl2*) [24]. On the other hand, Smg7 is linked to canonical ferroptotic effectors such as Hmox1, Ncoa4, Ftl1, Tfrc and Acsl4, thereby promoting hallmark ferroptotic processes including expansion of the labile iron pool, enhanced ferritinophagy and enrichment of polyunsaturated fatty acids in membrane phospholipids [25–28]. This regulatory pattern is consistent with the emerging roles of other NMD family members in ferroptosis [29]. In the multifactorial context of TBI, Smg7 thus co-activates mitochondrial apoptosis and ferroptosis, steering neurons toward Ferrapoptosis. Consequently, targeting Smg7 may afford broader neuroprotection than inhibiting any single death pathway alone.

Together with prior work on composite death modes such as PANoptosis and necroptotic apoptosis, our findings raise a broader question: might there be many more hybrid forms of cell death in vivo? The complexity of biological systems implies that signaling pathways rarely operate in isolation but instead form densely interconnected networks. Under certain conditions, converging lethal signals may drive the parallel engagement of multiple death programs. In this light, we view Ferrapoptosis as one example of a wider spectrum of potential hybrid entities and propose that concepts such as “hybrid” or “cross-over” regulated cell death may need to be incorporated into the current taxonomy. Systematically uncovering these composite death modes will deepen our understanding of cell fate control and open avenues for multi-target therapeutic strategies. We anticipate that future studies will increasingly focus on these crossroads within the cell death network, fostering the co-evolution of classification frameworks and precision intervention strategies.

## METHODS

### Study Design

Brain tissue for immunofluorescence (IF) was collected from mice subjected to mild, moderate, or severe TBI and the corresponding sham controls at acute, subacute, and chronic time points (n = 3 per severity and time point). An additional cohort of 12 mice was used to examine the effect of AAV-mediated Smg7 knockdown on histopathology. Animals were assigned to four groups: TBI + sh-NC (n = 3), sham + sh-NC (n = 3), TBI + sh-Smg7 (n = 3), and sham + sh-Smg7 (n = 3); cortical samples from these mice were processed for IF and TEM. For behavioral assessment, 18 mice were divided into three groups: TBI + sh-NC (n = 6), sham + sh-NC (n = 6), TBI + sh-Smg7 (n = 6). Behavioral testing was performed 30 days after surgery. The sample size and grouping for all in vivo and in vitro experiments are detailed in the corresponding methodological sections or indicated in the figure legends. To minimize bias, randomization and blinding were applied throughout: mice were randomly assigned to TBI or sham groups before surgery; investigators responsible for data collection, tissue processing, staining, and behavioral scoring were blinded to group allocation; and data analysts received only anonymized sample identifiers to ensure an unbiased interpretation of the results.

### Animals

C57BL/6J mice (6–8 weeks old, 20–23 g) were purchased from Beijing Huafukang Bioscience Co., Ltd. Animals were maintained in specific pathogen-free (SPF) facilities at Sichuan University under controlled conditions (22–25 °C, 50% relative humidity, 12-h light/dark cycle) with food and water available ad libitum. All procedures were approved by the Institutional Animal Care and Use Committee of Sichuan University (Approval No. K2021024) and conducted in accordance with relevant animal welfare guidelines. Both sexes were included in each experimental group, and mice were randomly assigned to experimental conditions.

### Single-nucleus RNA-seq analysis

Public single-nucleus RNA-seq data were obtained from mouse cortex at 24 h, 7 d and 6 months after TBI (GEO accession: GSE247339). Raw UMI count matrices were processed in Seurat (R v4.x). Nuclei with <200 or >5,000 detected genes or >10% mitochondrial transcripts were removed, and data were normalised and integrated using *SCTransform*. Principal component analysis was performed on highly variable genes, and a shared nearest-neighbour graph was used to compute a UMAP embedding. Pathway activity scores for ferroptosis and apoptosis were calculated at the single-nucleus level using AUCell with curated gene sets from MSigDB and recent literature. For each neuron, AUCell provided an AUC score for ferroptosis and apoptosis, which were projected onto the UMAP. Neurons with AUC values above the 70th percentile of the respective distribution were classified as “High”, and those at or below this threshold as “Low”. Combining the two pathways, we defined four phenotypes: ferroptosis-High-only, apoptosis-High-only, Double-High, and Double-Low. Changes in the proportions of these groups over time were summarised in bar and line plots. Differential expression and gene set enrichment analysis were then used to compare Double-High neurons with ferroptosis-High-only or apoptosis-High-only neurons and to identify pathways specifically enriched in Double-High cells.

### Human Tissue Samples and Ethics Statement

Human tissue samples were obtained from the Chengdu Public Security Bureau in compliance with the ethical principles outlined in the Declaration of Helsinki. The collection and use of human material were approved by the Ethics Committee of West China Hospital, Sichuan University (Approval No. 2022-153). Samples were collected from individuals who died from traumatic brain injury (TBI; n = 3) and from non-TBI controls (n = 3).

### CCI model

Mice were anesthetized with 4% isoflurane for induction and maintained with 1.5% isoflurane, then positioned in a stereotaxic frame. For moderate and severe TBI models, a circular craniotomy (∼4.5 mm in diameter) was made over the left parietal bone using a surgical drill, leaving the dura intact. A 3-mm diameter impactor tip was then used to deliver a controlled cortical impact through a sterile plastic film placed over the dura to minimize friction and contamination. For the mild TBI model, a direct cranial impact was applied without prior craniotomy. Impact parameters were set as follows: velocity 5.5 m/s and depth 3 mm for mild TBI, 5.0 m/s and 2 mm for moderate TBI, and 3.5 m/s and 1 mm for severe TBI [2, 30]. After impact, the scalp incision was closed with interrupted sutures and the wound was disinfected.

### Modified neurological severity score

Neurological function was assessed 6 h after injury using the modified Neurological Severity Score (mNSS). The test battery included assessments of motor function, sensory function, reflexes, and balance. One point was assigned for each failed task or absent reflex. Injury severity was classified according to the total mNSS as follows: 1–6, mild; 7–12, moderate; 13–18, severe TBI. Each animal was evaluated three times, and the mean score was used for subsequent statistical analysis.

### Cell culture and treatments

The HT22 mouse hippocampal neuronal cell line (RRID: CVCL_0321, kindly provided by Dr. Xiao Xiao of Sichuan University) was maintained in Dulbecco’s Modified Eagle Medium supplemented with 10% fetal bovine serum (FBS; Gibco, California, USA) and 1% penicillin-streptomycin (Gibco, California, USA), under standard culture conditions of 37°C and 5% CO□. A scratch assay was employed to simulate TBI in vitro. Briefly, a 100 μL pipette tip was used to create a uniform 5 × 5 grid of scratches on the confluent cell monolayer.

The culture medium was then carefully aspirated to remove detached cells and debris and replaced with fresh medium [30]. All chemical agents were initially dissolved in DMSO to prepare stock solutions, which were aliquoted and stored at –20 °C protected from light. Working solutions were prepared by diluting stocks in culture medium immediately before use. Final concentrations were as follows: 12 µM Erastin (Selleck Chemicals, Shanghai, China) to induce ferroptosis, 25 nM staurosporine (STS; MedChemExpress, New Jersey, USA) to induce apoptosis, and 27 mM sodium citrate to induce Ferrapoptosis. Control cells received an equivalent volume of DMSO without active compounds.

For experiments involving mitochondrial antioxidant pretreatment, cells were incubated with 500 nM MitoQ (Selleck Chemicals, Shanghai, China) for 24 h before subsequent injury paradigms and downstream analyses.

### Cell transfection

Smg7 siRNA (three independent oligos) and a negative control siRNA were synthesized by SeaStar Biotech (Jiangsu, China). A Smg7 overexpression plasmid (OE-Smg7) and the corresponding empty vector control (OE-NC) were constructed as described, and sequence information for all siRNA oligos and overexpression constructs is provided in Supplementary File 1. HT22 cells were transfected with 20 μM siRNA or with OE-Smg7/OE-NC plasmids using Lipofectamine 3000 (Invitrogen, Carlsbad, USA) in Opti-MEM serum-free medium according to the manufacturer’s instructions. After complex formation, transfection mixtures were added to cells maintained in complete DMEM and incubated for 48 h before downstream assays.

Smg7 siRNA sequences were as follows:

siRNA#1: sense, 5’-CGGCUUCUAUACUCAGUUATT-3’; antisense, 5’-UAACUGAGUAUAGAAGCCGTT-3’

siRNA#2: sense, 5’-CCAACAGCACAGUCUACAATT-3’; antisense, 5’-UUGUAGACUGUGCUGUUGGTT-3’

siRNA#3: sense, 5’-GGGUCAACAACGACGAAUATT-3’; antisense, 5’-UAUUCGUCGUUGUUGACCCTT-3’

### Transmission Electron Microscopy (TEM)

Ultrastructural analysis of cortical and hippocampal neurons was performed by TEM as previously described [31]. Mice were transcardially perfused with 0.1 M sodium cacodylate buffer containing 4% paraformaldehyde and 3% glutaraldehyde, and peri-lesional cortical tissue was post-fixed in 2% paraformaldehyde/2.5% glutaraldehyde. Samples were processed for resin embedding, ultrathin-sectioned (EM UC7, Leica, Germany), stained with uranyl acetate and lead citrate, and examined using a Hitachi transmission electron microscope (Hitachi, Japan).

### GSH/GSSG assays

Cellular glutathione was quantified using a GSH/GSSG Assay Kit (Beyotime, Shanghai, China) following the manufacturer’s instructions. Total glutathione (GSH + GSSG) and GSSG (after masking reduced GSH with 2-vinylpyridine) were measured spectrophotometrically at 412 nm (BioTek Synergy HT, CA, USA), and reduced GSH was calculated as total glutathione minus GSSG. Each condition was measured in six technical replicates.

### CCK-8 assays

Cell viability in 96-well plates was assessed using Cell Counting Kit-8 (CCK-8; Dojindo, Kumamoto, Japan). After addition of CCK-8 reagent to the culture medium, plates were incubated for 1 h at 37 °C, and absorbance at 450 nm was recorded using a microplate reader to determine relative viability.

### IF staining

For brain tissue, mice were perfused, and brains were fixed in 4% paraformaldehyde, cryoprotected, and coronally sectioned at 50 μm. Free-floating cortical sections were stored in PBS until staining. For cell staining, HT22 cells were grown on glass coverslips, exposed to the indicated treatments, and fixed in 4% paraformaldehyde for 15 min. Tissue sections and HT22 cells were processed using a TUNEL/IF workflow (Vazyme, Nanjing, China): PBS washes, permeabilization with 0.2% Triton X-100, and blocking with 5% bovine serum albumin. Samples were then incubated with primary antibodies against oxidized PRDX3-SO□/□ (Cat. #39806, Cayman Chemical) and NeuN (Cat. #94403, Cell Signaling Technology) overnight at 4 °C, followed by Alexa Fluor–conjugated secondary antibodies (Abcam) at room temperature. Apoptotic nuclei were labeled with a TUNEL kit (Beyotime), and nuclei were counterstained with DAPI. After final PBS washes, samples were mounted in antifade medium and imaged with a ZEISS LSM 980 confocal microscope.

### Annexin V-FITC / PI Staining

Apoptosis was assessed using an Annexin V–FITC/PI double-staining kit (KeyGen Biotech, Nanjing, China) according to the manufacturer’s instructions. Briefly, cells were harvested with EDTA-free trypsin, collected by centrifugation, and washed twice with PBS. Pellets were resuspended in 500 μL 1× Binding Buffer, followed by addition of 5 μL Annexin V–FITC and 5 μL propidium iodide (PI). After 5–10 min incubation at room temperature in the dark, stained cells were transferred to a 24-well plate and imaged by ZEISS LSM 980 confocal microscope.

### JC-1 Mitochondrial Membrane Potential Assay

Mitochondrial membrane potential was measured using a JC-1 Mitochondrial Membrane Potential Detection Kit (Cat. #abs50016, Absin, Shanghai, China) according to the manufacturer’s protocol. After removal of culture medium, adherent cells were incubated with JC-1-containing complete medium at 37 °C for 20 min, washed twice with prechilled 1× staining buffer, and equilibrated in fresh medium. Red (JC-1 aggregates) and green (monomers) fluorescence signals were acquired simultaneously on a ZEISS LSM 980 confocal microscope (Carl Zeiss AG).

### Lipid peroxidation assay

Lipid peroxidation was evaluated using the Image-iT™ Lipid Peroxidation Kit with BODIPY™ 581/591 C11 (Cat. #C10445, Invitrogen, Thermo Fisher Scientific, USA) according to the manufacturer’s instructions. Treated HT22 cells were incubated with 10 μM BODIPY 581/591 C11 in fresh complete medium at 37 °C for 30 min, washed three times with PBS, and imaged immediately by epifluorescence or confocal microscopy (red channel Ex/Em 581/591 nm; green channel Ex/Em 488/510 nm). Lipid peroxidation was quantified as the red/green fluorescence ratio. For brain tissue, 10-μm frozen sections prepared from 4% paraformaldehyde-fixed, 30% sucrose-cryoprotected samples were rehydrated in PBS, incubated with 10 μM probe for 30 min, washed, and imaged by confocal microscopy for red/green ratio analysis.

### FerroOrange Staining

Intracellular Fe²□ was detected using the FerroOrange fluorescent probe (Cat. #F374, Dojindo Laboratories, Kumamoto, Japan) following the manufacturer’s instructions. After the indicated treatments, HT22 cells were washed twice with PBS and incubated with 1 μM FerroOrange working solution at 37 °C for 30 min. Excess probe was removed by PBS washes, and fluorescence images were acquired using ZEISS LSM 980 confocal microscope.

### Bulk RNA-seq sample preparation and data analysis

Mouse hippocampal HT22 cells were cultured, transfected with siNC or siSmg7, and subjected to scratch injury as described above. Cells from different groups were collected for bulk RNA-seq, and total RNA was isolated and used for standard library preparation and Illumina sequencing by a commercial service provider. Raw count matrices were imported into R, and genes with zero counts across all samples were removed. Ensembl gene identifiers were stripped of version numbers and mapped to mouse gene symbols using org.Mm.eg.db, with unresolved IDs complemented by biomaRt queries. Counts were normalized to log□ counts-per-million using edgeR, and differential expression between groups was assessed with limma using a linear model ∼ group and empirical Bayes moderation. Significantly regulated genes were defined by Benjamini–Hochberg adjusted P < 0.05 and |log□ fold-change| > 1, and visualized as heatmaps and volcano plots. Ferroptosis- and apoptosis-related genes were extracted from KEGG pathways “Ferroptosis” (mmu04216) and “Apoptosis” (mmu04210) for pathway-focused heatmaps and boxplots, and functional enrichment of differentially expressed genes was performed with clusterProfiler for Gene Ontology and KEGG analyses.

## REFERENCES

1. Maas AIR, Menon DK, Adelson PD, Andelic N, Bell MJ, Belli A, et al. Traumatic brain injury: integrated approaches to improve prevention, clinical care, and research. Lancet Neurol. 2017;16(12):987–1048. Epub 20171106. doi: 10.1016/S1474-4422(17)30371-X. PubMed PMID: 29122524.

2. Robinson CP. Moderate and Severe Traumatic Brain Injury. Continuum (Minneap Minn). 2021;27(5):1278–300. doi: 10.1212/CON.0000000000001036. PubMed PMID: 34618760.

3. Moujalled D, Strasser A, Liddell JR. Molecular mechanisms of cell death in neurological diseases. Cell Death Differ. 2021;28(7):2029–44. Epub 20210607. doi: 10.1038/s41418-021-00814-y. PubMed PMID: 34099897; PubMed Central PMCID: PMCPMC8257776.

4. Eskander G, Abdelhamid SG, Wahdan SA, Radwan SM. Insights on the crosstalk among different cell death mechanisms. Cell Death Discov. 2025;11(1):56. Epub 20250210. doi: 10.1038/s41420-025-02328-9. PubMed PMID: 39929794; PubMed Central PMCID: PMCPMC11811070.

5. Jiang L, Kon N, Li T, Wang SJ, Su T, Hibshoosh H, et al. Ferroptosis as a p53-mediated activity during tumour suppression. Nature. 2015;520(7545):57–62. Epub 20150318. doi: 10.1038/nature14344. PubMed PMID: 25799988; PubMed Central PMCID: PMCPMC4455927.

6. Neitemeier S, Jelinek A, Laino V, Hoffmann L, Eisenbach I, Eying R, et al. BID links ferroptosis to mitochondrial cell death pathways. Redox Biol. 2017;12:558–70. Epub 20170309. doi: 10.1016/j.redox.2017.03.007. PubMed PMID: 28384611; PubMed Central PMCID: PMCPMC5382034.

7. Qiu Y, Huther JA, Wank B, Rath A, Tykwe R, Aldrovandi M, et al. Interplay of ferroptotic and apoptotic cell death and its modulation by BH3-mimetics. Cell Death Differ. 2025;32(11):1970–85. Epub 20250429. doi: 10.1038/s41418-025-01514-7. PubMed PMID: 40301648; PubMed Central PMCID: PMCPMC12572382.

8. Cui S, Ghai A, Deng Y, Li S, Zhang R, Egbulefu C, et al. Identification of hyperoxidized PRDX3 as a ferroptosis marker reveals ferroptotic damage in chronic liver diseases. Mol Cell. 2023;83(21):3931–9 e5. Epub 20231019. doi: 10.1016/j.molcel.2023.09.025. PubMed PMID: 37863053; PubMed Central PMCID: PMCPMC10841858.

9. Wu Y, Jia C, Liu W, Zhan W, Chen Y, Lu J, et al. Sodium citrate targeting Ca(2+)/CAMKK2 pathway exhibits anti-tumor activity through inducing apoptosis and ferroptosis in ovarian cancer. J Adv Res. 2024;65:89–104. Epub 20240507. doi: 10.1016/j.jare.2024.04.033. PubMed PMID: 38724006; PubMed Central PMCID: PMCPMC11518946.

10. Li D, Huang S, Zhu J, Hu T, Han Z, Zhang S, et al. Exosomes from MiR-21-5p-Increased Neurons Play a Role in Neuroprotection by Suppressing Rab11a-Mediated Neuronal Autophagy In Vitro After Traumatic Brain Injury. Med Sci Monit. 2019;25:1871–85. Epub 20190312. doi: 10.12659/MSM.915727. PubMed PMID: 30860987; PubMed Central PMCID: PMCPMC6423733.

11. Usuki F, Yamashita A, Fujimura M. Environmental stresses suppress nonsense-mediated mRNA decay (NMD) and affect cells by stabilizing NMD-targeted gene expression. Sci Rep. 2019;9(1):1279. Epub 20190204. doi: 10.1038/s41598-018-38015-2. PubMed PMID: 30718659; PubMed Central PMCID: PMCPMC6362056.

12. Luo H, Cowen L, Yu G, Jiang W, Tang Y. SMG7 is a critical regulator of p53 stability and function in DNA damage stress response. Cell Discov. 2016;2:15042. Epub 20160119. doi: 10.1038/celldisc.2015.42. PubMed PMID: 27462439; PubMed Central PMCID: PMCPMC4860962.

13. Goetz AE, Wilkinson M. Stress and the nonsense-mediated RNA decay pathway. Cell Mol Life Sci. 2017;74(19):3509–31. Epub 20170513. doi: 10.1007/s00018-017-2537-6. PubMed PMID: 28503708; PubMed Central PMCID: PMCPMC5683946.

14. Zhou SY, Cui GZ, Yan XL, Wang X, Qu Y, Guo ZN, et al. Mechanism of Ferroptosis and Its Relationships With Other Types of Programmed Cell Death: Insights for Potential Interventions After Intracerebral Hemorrhage. Front Neurosci. 2020;14:589042. Epub 20201113. doi: 10.3389/fnins.2020.589042. PubMed PMID: 33281547; PubMed Central PMCID: PMCPMC7691292.

15. Xiong C, Ling H, Hao Q, Zhou X. Cuproptosis: p53-regulated metabolic cell death? Cell Death Differ. 2023;30(4):876–84. Epub 20230208. doi: 10.1038/s41418-023-01125-0. PubMed PMID: 36755067; PubMed Central PMCID: PMCPMC10070433.

16. Bramlett HM, Dietrich WD. Long-Term Consequences of Traumatic Brain Injury: Current Status of Potential Mechanisms of Injury and Neurological Outcomes. J Neurotrauma. 2015;32(23):1834–48. Epub 20141219. doi: 10.1089/neu.2014.3352. PubMed PMID: 25158206; PubMed Central PMCID: PMCPMC4677116.

17. Huang Q, Zhang H, Chen S, Wang Y, Zhou J. Ferroptosis in central nervous system injuries: molecular mechanisms, diagnostic approaches, and therapeutic strategies. Front Cell Neurosci. 2025;19:1593963. Epub 20250722. doi: 10.3389/fncel.2025.1593963. PubMed PMID: 40766185; PubMed Central PMCID: PMCPMC12321900.

18. Verma M, Lizama BN, Chu CT. Excitotoxicity, calcium and mitochondria: a triad in synaptic neurodegeneration. Transl Neurodegener. 2022;11(1):3. Epub 20220125. doi: 10.1186/s40035-021-00278-7. PubMed PMID: 35078537; PubMed Central PMCID: PMCPMC8788129.

19. Li QS, Jia YJ. Ferroptosis: a critical player and potential therapeutic target in traumatic brain injury and spinal cord injury. Neural Regen Res. 2023;18(3):506–12. doi: 10.4103/1673-5374.350187. PubMed PMID: 36018155; PubMed Central PMCID: PMCPMC9727428.

20. Tang S, Gao P, Chen H, Zhou X, Ou Y, He Y. The Role of Iron, Its Metabolism and Ferroptosis in Traumatic Brain Injury. Front Cell Neurosci. 2020;14:590789. Epub 20200925. doi: 10.3389/fncel.2020.590789. PubMed PMID: 33100976; PubMed Central PMCID: PMCPMC7545318.

21. Hu X, Xu Y, Zhang H, Li Y, Wang X, Xu C, et al. Role of necroptosis in traumatic brain and spinal cord injuries. J Adv Res. 2022;40:125–34. Epub 20211222. doi: 10.1016/j.jare.2021.12.002. PubMed PMID: 36100321; PubMed Central PMCID: PMCPMC9481937.

22. Yang L, Kraft VAN, Pfeiffer S, Merl-Pham J, Bao X, An Y, et al. Nonsense-mediated decay factor SMG7 sensitizes cells to TNFalpha-induced apoptosis via CYLD tumor suppressor and the noncoding oncogene Pvt1. Mol Oncol. 2020;14(10):2420–35. Epub 20200713. doi: 10.1002/1878-0261.12754. PubMed PMID: 32602581; PubMed Central PMCID: PMCPMC7530794.

23. Bratton SB, Salvesen GS. Regulation of the Apaf-1-caspase-9 apoptosome. J Cell Sci. 2010;123(Pt 19):3209–14. doi: 10.1242/jcs.073643. PubMed PMID: 20844150; PubMed Central PMCID: PMCPMC2939798.

24. Li H, Zhu H, Xu CJ, Yuan J. Cleavage of BID by caspase 8 mediates the mitochondrial damage in the Fas pathway of apoptosis. Cell. 1998;94(4):491–501. doi: 10.1016/s0092-8674(00)81590-1. PubMed PMID: 9727492.

25. Fang X, Cai Z, Wang H, Han D, Cheng Q, Zhang P, et al. Loss of Cardiac Ferritin H Facilitates Cardiomyopathy via Slc7a11-Mediated Ferroptosis. Circ Res. 2020;127(4):486–501. Epub 20200430. doi: 10.1161/CIRCRESAHA.120.316509. PubMed PMID: 32349646.

26. Yang X, Ding Y, Sun L, Shi M, Zhang P, Huang Z, et al. Ferritin light chain deficiency-induced ferroptosis is involved in preeclampsia pathophysiology by disturbing uterine spiral artery remodelling. Redox Biol. 2022;58:102555. Epub 20221124. doi: 10.1016/j.redox.2022.102555. PubMed PMID: 36446230; PubMed Central PMCID: PMCPMC9706170.

27. Chen X, Yu C, Kang R, Tang D. Iron Metabolism in Ferroptosis. Front Cell Dev Biol. 2020;8:590226. Epub 20201007. doi: 10.3389/fcell.2020.590226. PubMed PMID: 33117818; PubMed Central PMCID: PMCPMC7575751.

28. Li A, Li Y, Li X, Tang C, Yang Y, Li N, et al. Ferritin light chain as a potential biomarker for the prognosis of liver hepatocellular carcinoma. Heliyon. 2024;10(16):e36040. Epub 20240809. doi: 10.1016/j.heliyon.2024.e36040. PubMed PMID: 39224384; PubMed Central PMCID: PMCPMC11367121.

29. Han L, Bai L, Fang X, Liu J, Kang R, Zhou D, et al. SMG9 drives ferroptosis by directly inhibiting GPX4 degradation. Biochem Biophys Res Commun. 2021;567:92–8. Epub 20210616. doi: 10.1016/j.bbrc.2021.06.038. PubMed PMID: 34146907.

30. Manrui L, Xu Y, Liu J, Zhang X, Yuan R, Sun Y, et al. Aminophylline targets miR-128-3p/Slc7a11 axis to attenuate neuronal ferroptosis after traumatic brain injury. Cell Mol Life Sci. 2025;82(1):87. Epub 20250222. doi: 10.1007/s00018-025-05601-3. PubMed PMID: 39985589; PubMed Central PMCID: PMCPMC11846823.

31. Yang Q, Zhang L, Li M, Xu Y, Chen X, Yuan R, et al. Single-nucleus transcriptomic mapping uncovers targets for traumatic brain injury. Genome Res. 2023;33(10):1818–32. Epub 20230920. doi: 10.1101/gr.277881.123. PubMed PMID: 37730437; PubMed Central PMCID: PMCPMC10691476.

