## Supplemental Figure for "Ferrapoptosis: Discovery of a Hybrid Programmed Cell Death"

**List of Supplementary Materials**

Fig. S1 to Fig. S2

Supplementary File 1: OE-Smg7 sequencing results

Supplementary File 2: Viral vector sequencing results

**Figure S1**

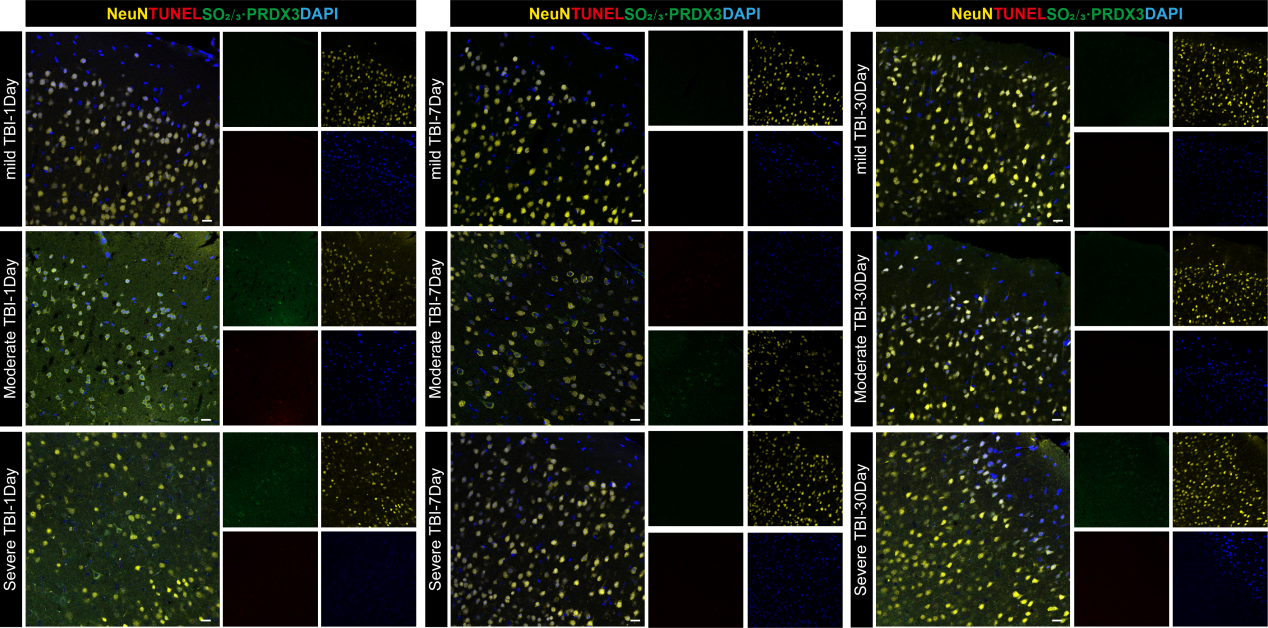

**Figure S1. Contralateral cortical immunofluorescence for NeuN, TUNEL and PRDX3-SO₂/₃ after TBI.**

Representative confocal images of the contralateral cortex from mice subjected to mild, moderate or severe TBI at 1, 7 and 30 days, stained for NeuN (yellow), TUNEL (red), oxidized PRDX3-SO₂/₃ (green) and DAPI (blue). n = 3 mice per group. Scale bars, as indicated.

**Figure S2**

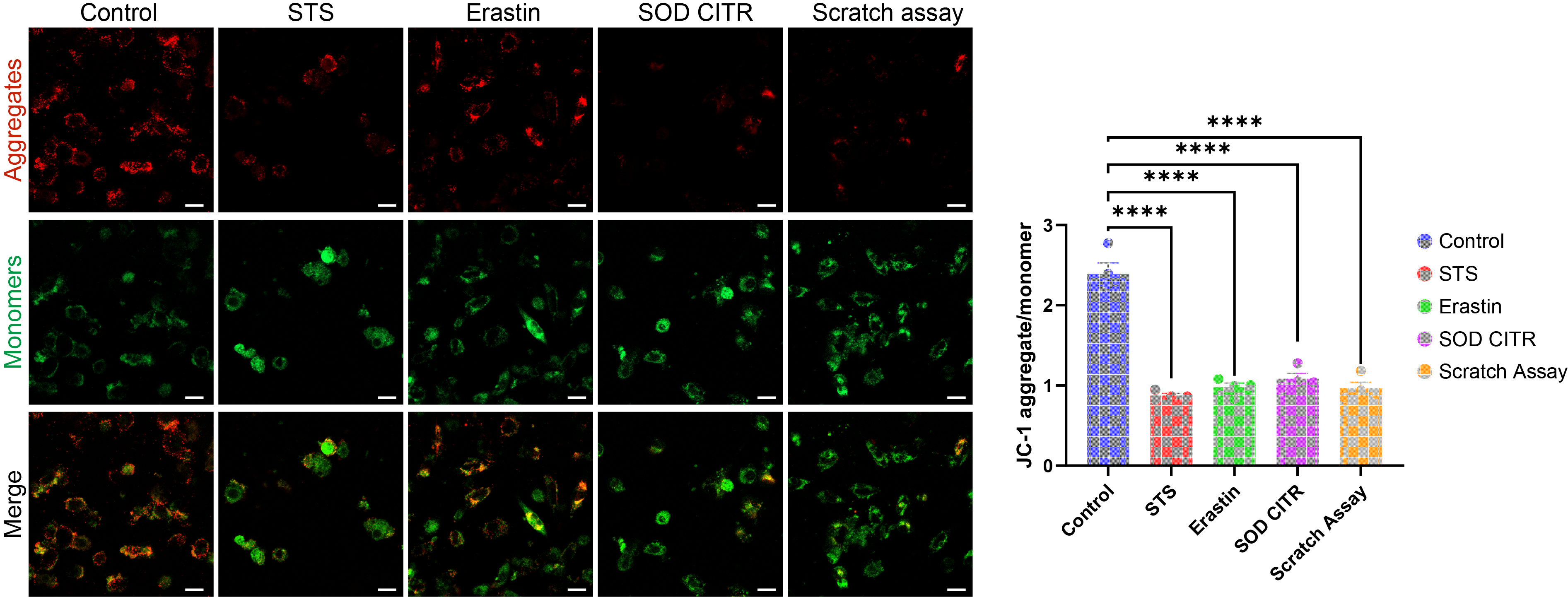

**Figure S2. JC-1 assessment of mitochondrial membrane potential in HT22 cells.**
Representative JC-1 staining images (aggregates, red; monomers, green; merged) of HT22 cells 12 h after treatment with vehicle (Control), STS, Erastin, sodium citrate (SOD CITR) or scratch injury, and corresponding quantification of the JC-1 aggregate/monomer fluorescence ratio. Cells were stained with JC-1 according to the manufacturer’s protocol and imaged by confocal microscopy. Data are presented as mean ± SEM (n = 3 independent cultures per group) and analyzed by one-way ANOVA followed by Tukey’s post hoc test; ns, not significant; ****P < 0.0001. Scale bars=20μm.

**Supplementary File 1: OE-Smg7 sequencing results**

CAGGTCCAACTGCACCTCGGTTCTGCTAGCCGCCACCATGAGCCTGCAGAGCGCGCAGTATCTCCGGCAGGCAGAAGTCCTGAAGGCTGAAATGACAGATTCTAAGCTGGGTCCGGCTGAGGTCTGGACATCCAGGCAAGCTCTGCAGGACTTGTACCAGAAAATGCTAGTTACTGATCTGGAATACGCTTTAGACAAGAAAGTAGAACAGGATCTCTGGAATCATGCCTTTAAGAATCAGATCACAACACTGCAAGGCCAAGCCAAGAATCGAGCAAACCCAAATCGAAGTGAAGTCCAGGCAAACCTTTCTCTGTTCCTAGAGGCAGCTAGCGGCTTCTATACTCAGTTATTACAGGAACTGTGTACGGTGTTTAATGTAGATTTGCCATGCCGTGTGAAGTCTTCCCAGTTGGGAATTATTAGCAATAAACAGACGCACAGCAGCACCATAGTGAAGCCACAGTCTAGCTCCTGCTCCTACATTTGCCAACACTGCCTCGTCCACCTTGGAGACATTGCTCGATACAGAAATCAGACTAGCCAGGCAGAGTCCTACTATAGGCATGCAGCTCAGCTTGTCCCCTCCAATGGTCAGCCTTACAATCAGTTGGCTATCTTAGCTTCTTCCAAAGGAGACCACCTGACCACAATTTTCTACTACTGCAGAAGCATTGCTGTGAAATTCCCTTTCCCAGCTGCCTCCACTAATCTACAAAAAGCACTTTCTAAAGCACTGGAAAGCCGGGATGAGTTGAAAACCAAGTGGGGTGTTTCTGACTTCATCAAGGCCTTTATTAAATTCCACGGTCATGTATACCTGAGTAAGAGTTTGGAAAAGCTGAGCCCTCTTCGAGAGAAGTTGGAAGAACAGTTTAAGAGACTGCTGTTCCAAAAGGCTTTCAACTCTCAACAGTTAGTTCATGTCACTGTCATTAACCTGTTTCAACTTCATCATCTTCGTGACTTTAGCAATGAAACAGAGCAACACAGTTACAGCCAAGATGAACAGCTGTGTTGGACACAGTTGCTGGCCCTCTTTATGTCTTTTCTCGGCATCCTGTGCAAATGTCCTCTCCAGAATGATTCTCAGGAGTCCAACAATGCCTACCCCCTTCCTGCAGTCAAGGTCTCCATGGACTGGCTAAGACTCCGACCTAGAGTCTTTCAAGAAGCAGTGGTGGATGAAAGACAGTACATTTGGCCCTGGCTAATTTCTCTTCTAAATAGTTTCCATCCCCGTGAAGATGATCTCTCAAATACTAATGCCACACCACTTCCAGAGGAGTTTGAATTACAAGGATTCTTGGCTTTAAGACCTTCTTTCAGGAACTTGGATTTTTCCAAAGGCCATCAGGGTATTACAGGAGACAAAGAGGGTCAACAACGACGAATACGACAGCAGCGTTTGATCTCCATAGGGAAATGGATTGCTGATAACCAGCCACGGCTGATTCAGTGTGAAAATGAGGTAGGGAAATTGTTATTTATCACAGAAATCCCAGAATTAATACTAGAAGACCCCAGTGAAGCCAAAGAGAACCTCATCCTACAAGAAACATCTGTGGTAGAGTCACTGGCTACGGATGGGAGCCCAGGACTGAAATCAGTGCTGTCTACAGGCCGAAATCCAAGCAACAGCTGTGACTCAGGAGAGAAACCAGTGGTCACCTTTAAAGAGAACATTAAGCCACGAGAAGTGAACCAAGGAAGAAGCTTTCCTCCCAAAGAGGTAAAATCCCAGACAGAACTAAGAAAGACTCCAGTGTCTGAAGCCAGGAAAACGCCTGTCACTCAAACTCCAAGTCAAACGAGTAATTCTCAGTTCATCCCCATCCATCACCCTGGAGCCTTCCCTCCTCTTCCCAGCCGACCAGGATTCCCGCCCCCAACATATGTTATCCCCCCTCCTGTGGCATTTTCTATGGGCTCAGGTTACACCTTCCCAGCTGGTGTTTCTGTCCCAGGAACCTTTCTTCAGTCTACAGCTCACTCTCCAGCAGGAAACCAGGTGCAAGCTGGGAAACAGTCCCACATTCCTTACAGCCAGCAACGGCCCTCTGGACCAGGGCCAATGAACCAGGGACCTCAACAGTCACAGCCACCTTCCCAGCCACCCCTTACATCTTTACCAGCTCAGCCAACAGCACAGTCTACAAGCCAATTGCAGGTTCAAGCTCTAGCTCAGCAACAGCAATCCCCTACAAAAGTCATACCAGCTTTGGGGAAAAGCCCGCCTCACCACTCTGGATTCCAGCAGTATCAACAGGCAGATGCCTCCAAACAGCTGTGGAATCCCCCTCAGGTTCAAAGCCCACTAGGGAAAATTATGCCTGTGAAACAGTCCTACTACCTTCAGACCCAAGACCCTATAAAACTGTTTGAGCCGTCATTGCAACCTCCTGTAATACAGCAGCAGCCTCTAGAGAAAAAAATGAAGCCTTTCCCCATGGAGCCATATAACCATAATCCCTCAGAAGTCAAGGTCCCAGAGTTCTACTGGGATTCTTCCTACAGCATGGCCGATAACAGAGCAGTAATGGCTCAGCAACCAAATATGGACCGCAGGAGCAAACGGTCACCTGGAGTCTTCCGTCCAGAACAGGATCCTGTGCCCAGGATGCCATTTGAGGACCCCAAGAGCTCCCCTCTGCTTCCTCCGGACCTGTTAAAGAGTCTGGCTGCCTTGGAGGAAGAGGAAGAGCTGATCTTTTCTAACCCTCCTGATCTTTACCCAGCTCTGCTGGGTCCTCTCGCCTCTCTTCCTGGACGAAGCCTCTTTAAATCCTTATTGGAGAAGCCTTCTGAGCTCATGTCACATTCATCTTCTTTCCTGTCCCTCACGGGGTTCTCTGTCAATCAGGAAAGATATCCAAACAGCAGTATGTTCAATGAAGTGTATGGAAAAAACCTGACAACCAGCTCCAAGGCAGAACTGAACCCCTCAGTCGCCTCCCAGGAAACATCACTGTACTCCCTTTTTGAAGGGACCCCATGGTCTCCATCACTTCCTGCCAGTTCAGATCATTCGACACCAGCCAGCCAGTCTCCTCATTCCTCTAACCCAAGCAGCCTGCCCAGTTCTCCTCCAACGCATAACCATAATTCTGCTCCATTCTCTAATTTTGGACCCATTGGGACTCCAGATAACAGGGATAGGCGGCCTGCAGATAGGTGGAAAACTGATAAACCAGCCATGGGTGGTTTTGGTGTTGATTATCTCTCAGCAACGTCATCATCTGAGAGCAGTTGGCATCAGGCTAGCACTCCGAGTGGCACCTGGACAGGCCACGGCCCCTCCATGGAGGATTCCTCTGCTGTCCTCATGGAAAGCCTAAAGTCTATCTGGTCCAGTTCCATGATGCATCCTGGGCCCTCCGCTCTGGAGCAGTTGTTAATGCAGCAGAAGCAGAAACAGCAGCGGGGACAAGGTGCCATGAACCCTCCACACACGGGCCCTCTAGACGCCACCATGGTGAGCAA

**Supplementary File 2: Viral vector sequencing results**

>P25F2555-1-RT105F_F11.ab1GGCTAGGCGACTCTTCAGTCCGCCATGCCCGAAGGCTACGTCCAGGAGCGCACCATCTTCTTCAAGGACGACGGCACTACAGACCCGCGCCGAGGTGAAGTTCGAGGGCGACACCCTGGTGAACCGCATCGAGCTGAAGGGCATCGACTTCAAGGAGGACGGCAACATCCTGGGGCACAAGCTGGAGTACAACTACAACAGCCACAACGTCTATATCATGGCCGACAAGCAGAAGAACGGCATCAAGGTGAACTTCAAGATCCGCCACAACATCGAGGACGGCAGCGTGCAGCTCGCCGACCACTACCAGCAGAACACCCCCATCGGCGACGGCCCCGTGCTGCTGCCCGACAACCACTACCTGAGCACCCAGTCCGCCCTGAGCAAAGACCCCAACGAGAAGCGCGATCACATGGTCCTGCTGGAGTTCGTGACCGCCGCCGGGATCACTCTCGGCATGGACGAGCTGTACAAGGCTAGCTAACTGGAGGCTTGCTGAAGGCTGTATGCTGTATTCGTCGTTGTTGACCCTCGTTTTGGCCACTGACTGACGAGGGTCAAACGACGAATACAGGACACAAGGCCTGTTACTAGCACTCACATGGAACAAATGGCCCAAGCTTTAAACCGGTTATCGATAATCAACCTCTGGATTACAAAATTTGTGAAAGATTGACTGGTATTCTTAACTATGTTGCTCCTTTTACGCTATGTGGATACGCTGCTTTAATGCCTTTGTATCATGCTATTGCTTCCCGTATGGCTTTCATTTTCTCCTCCTTGTATAAATCCTGGTTTGCTGTCTCTTTATGAGGAGTTGTGGCCCGTTGTCAGGCAACGTGCGTGTGTGCACTGTGTTTGCTGACGCAACCCCCACTGTTGGGGCATTGCCACCACCTGTCAGCTCCTTTCCGGAACTTTCGCTTTCCCCCCTCCCTATTGCCACGGCGGAACTCATCGCCGCCTGCCTTGGCC

P25F2555 GACGAGCTGTACAAGGCTAGCTAACTGGAGGCTTGCTGAAGGCTGTATGCTGTATTCGTCGTTGTTGACCCTCGTTTTGGCCACTGACTGACGAGGGTCAAACGACGAATACAGGACACAAGGCCTGTTACTAGCACTCACATGGAACAAATGGCCCAAGCTTTAAACCGGTTATCGA
